# Large body size variation is linked to low communication success in tandem running ants

**DOI:** 10.1101/789834

**Authors:** Wagner Thomas, Bachenberg Lena, Glaser Simone, Oikonomou Avgousta, Linn Melissa, Grüter Christoph

## Abstract

Diversity in animal groups is often assumed to increase group performance. In insect colonies, genetic, behavioral and morphological variation among workers can improve colony functioning and resilience. However, it has been hypothesized that during communication processes, differences between workers, e.g. in body size, could also have negative effects. Tandem running is a common recruitment strategy in ants and allows a leader to guide a nestmate follower to resources. A substantial proportion of tandem runs fail because leader and follower loose contact. Using the ant *Temnothorax nylanderi* as a model system, we tested the hypothesis that tandem running success is impaired if leader and follower differ in size. Indeed, we found that the success rate of tandem pairs drops considerably as size variation increases: only ~7% of tandem runs were successful when the leader-follower size difference exceeded 10%, whereas 80% of tandem runs were successful when ants differed less than 5% in body length. One possible explanation is that ant size is linked to the preferred walking speed. Ants did not choose partners of similar size, but extranidal workers were larger than intranidal workers, which could reduce recruitment mistakes because it reduced the chance that very large and very small ants perform tandem runs together. Our results suggest that phenotypic differences between interacting workers can have negative effects on the efficiency of communication processes. Whether phenotypic variation has positive or negative effects is likely to depend on the task and the phenotypic trait that shows variation.

## Introduction

Social groups consist of individuals that differ from each other in a number of ways. For instance, people working in a company may differ in experience, training, gender, ethnicity or skills and this diversity can affect group performance and success. Scientists interested in organizational theory have found that group diversity often has positive effects on group performance, most likely because diverse groups possess a broader range of knowledge, experience, personality, skills and abilities (Horwitz and Horwitz 2007; van Knippenberg and Schippers 2007). For instance, more diverse scientific collaborations produce publications of greater impact (Freeman and Huang 2015; Alshebli et al. 2018), more ethnically diverse markets show lower risks of price bubbles (Levine et al. 2014) and companies with greater gender diversity in corporate leadership positions are more successful (Noland et al. 2016). As a result, organizations may compose teams to incorporate differences in functional, ethnic or educational background (van Knippenberg and Schippers 2007). However, group diversity can also have adverse effects on consensus decision-making, group functioning and increase intra-group conflicts (Horwitz and Horwitz 2007; van Knippenberg and Schippers 2007). One reason is that individuals might prefer to interact with more similar individuals.

Insect colonies consist of many individuals that may appear similar or even identical to the casual observer, but closer examination readily reveals that the members of a colony differ in many ways, including their morphology, behavior, experience or genetic background (Wilson 1971; Oster and Wilson 1978; Hölldobler and Wilson 2009). Several studies have found that differences among workers promote colony success. For example, colonies that show greater behavioral variation, e.g. because they are genetically more diverse, have been shown to collect food more successfully, respond better to environmental perturbations or produce more brood (Jones et al. 2004; Mattila and Seeley 2007; Oldroyd and Fewell 2007; Modlmeier and Foitzik 2011). Workers of many species also differ in their morphology and this morphological variation is closely tied to division of labor in many species (ants: Hölldobler and Wilson 2009; bumblebees: Goulson et al. 2002; stingless bees: Grüter et al. 2017a, Baudier et al. 2019; termites: Tian & Zhou 2014). Having different worker types for different tasks is likely to increase group performance because different worker types are more efficient at performing particular tasks (Oster and Wilson 1978; Powell and Franks 2005; Mertl and Traniello 2009; Grüter et al. 2012, 2017b; Powell 2016). Even in species with gradual variation, *i.e.* without distinct morphological castes, colonies with a larger worker size range often seem more successful (Porter and Tschinkel 1985; Beshers and Traniello 1994; Billick 2002; Billick and Carter 2007, but see Jandt and Dornhaus 2014; Colin et al. 2017). In most cases, this intra-colonial variation is an example of phenotypic plasticity, where the phenotypic differences are generated by variation in environmental factors (e.g. food quantity, temperature), rather than differences in genotype (Oster and Wilson 1978; Sumner et al. 2006; Segers et al. 2015; Molet et al. 2017).

However, there could also be circumstances where differences among cooperating workers have negative effects on group performance. Waddington et al. (1986) and Waddington (1989) argued that worker size variation reduces the efficiency of communication in social bees. If signal producers and receivers differ in morphology (and their sensory systems), the communication of misinformation or impaired information transfer could become more likely. Consistent with this hypothesis is the finding that stingless bees with more sophisticated communication show lower within colony size variation (Waddington et al. 1986) and honeybee (*Apis mellifera*) colonies with larger body size variation collected less nectar (Waddington 1989). The latter study also found that honeybee dancers tended to interact with bees of similar size. Waddington et al. (1986) argue that negative effects of size differences on communication could explain why colonies of the highly eusocial honeybees and stingless bees, which often use sophisticated communication during foraging, are less morphologically variable than the primitively eusocial bumblebees, which use simpler methods of communication (Dornhaus and Chittka 1999). However, the potential disadvantages of worker size variation for communication processes in insect societies has received little attention and, as far as we know, we still lack evidence that body size differences among interacting individuals indeed affects communication efficiency.

A recruitment behaviour that relies on communication and that could be negatively affected by body size differences is tandem running, which is relatively common in ant species with small colony sizes (so far described in ~40 species) (Beckers et al. 1989; Franklin 2014; Grüter et al. 2018). During a tandem run, a leader with information about the location of a resource slowly guides a nestmate to a food source or a nest site. Contact between the leader and the follower is maintained by frequent physical interactions and short-range pheromones (Möglich et al. 1974; Basari et al. 2014*a*). This behavior has been considered a case of animal teaching because leaders actively facilitate learning in a nestmate while also incurring costs (Franks and Richardson 2006). Contact loss is common during tandem runs, but break-ups are often prevented by leaders waiting for their partner while the latter searches for her leader (Franks and Richardson 2006; Richardson et al. 2007). Nonetheless, a substantial proportion of tandem runs break up before reaching the goal (e.g. ~19% in *Temnothorax nylanderi*, Glaser and Grüter 2018; ~23% in *Pachycondyla harpax*, Grüter et al. 2018; ~50% in *Cardiocondyla venustula*, Wilson 1959; up to 70% in *Temnothorax rugatulus*, Pratt 2008). Break-ups are costly in terms of time and could leave lost followers in dangerous areas.

We studied tandem running during colony emigrations in *Temnothorax nylanderi*, a species with moderate size variation (Molet et al. 2017), and tested the hypothesis that body size differences between leaders and followers affects the efficiency of recruitment. Specifically, we tested the prediction that size differences between interacting partners have a negative effect on the success rate. To obtain a better understanding of body size variation in this population, we also quantified the body size distribution in several colonies and tested whether extranidal workers (potential scouts for nest-sites or food) are larger than intranidal workers, as was found in two other species from the same genus (Herbers and Cunningham 1983; Westling et al. 2014). Finally, we measured the relationship between body size and the walking speed of an ant.

## Methods

### Study site and species

*Temnothorax nylanderi* colonies were collected from acorns and decaying branches in the Lenneberg forest (50°00’44.2 N, 8°10’57.8 E) near Mainz in Germany in 2015 (part 1), 2016 (part 2), 2017 (part 4) and 2018 (part 3). Back in the laboratory, colonies were kept in nests made up of two microscope slides (50 mm × 10 mm × 3 mm) and, between the slides, an acrylic glass slide containing an oval cavity that created a living space. This nest was placed in a larger plastic box (100 mm × 100 mm × 30 mm) with paraffin oil-coated walls that prevented ants from escaping. Colonies had a reproductive queen and brood and were kept in a climate chamber at 22°C with a 12:12 h light/dark cycle. Colonies were fed twice a week with honey and a cricket and were provided with water *ad libitum*. *Temnothorax* colonies can occasionally contain queen-worker “intercastes”, but they are very rare (Okada et al. 2013) and none were found in our colonies.

#### Part 1: Measuring the body size distribution

All workers from four recently collected colonies (colony size: 59-76 workers) were individually transferred into a petri dish covered with graph paper, which served as a scale. Each ant was photographed three times with a Nikon D7000 camera (AF-S Micro Nikkor 105 mm lens), mounted on a tripod and at a constant distance of approx. 30 cm above the petri dish. We measured total body length and head width of ants using ImageJ 1.46 and averaged the values from the three photos. Body length and head width were highly correlated (r = 0.84, N = 262, Pearson correlation: *p* < 0.0001), but body length is used as our measure of body size in this study because it is easier to measure due to the small size of the ants.

#### Part 2: Body size and walking speed

We collected 3–4 large (2.7–3.3 mm body length) and 3–4 small (2.2–2.69 mm) workers from each of eight colonies as they were walking outside their artificial nest and put them in small groups in a plastic arena (17.8 × 11.8 × 4.7 cm). The walls were coated with Fluon to prevent ants from escaping. The floor was covered with graph paper that allowed us to measure walking speed. Body size was measured as described above. After a 10 min acclimatization period, ants were filmed (Canon Legria HF R706) from above (30 cm distance) for 10 min as they were exploring the arena. The walking speed (cm/sec) of ants was determined using the object detection and tracking software *AnTracks* (www.antracks.org). Three uninterrupted 20 sec sequences (*i.e.* periods when ants did not interact with another ant, stand still or walk along a wall) were averaged for each ant.

#### Part 3: Body size and tandem running

We used 22 colonies (range of 56-100 workers). One day before an emigration, nests were placed in an emigration arena (31.2 × 22.3 × 4.7 cm). The walls were again coated with Fluon and the floor was covered with graph paper. On a test day, a new empty nest was placed in the emigration arena, 20 cm from the original nest. To motivate the colony to move to this new nest, the old nest was destroyed by gently removing the lid (see e.g. Mallon and Franks 2000; Pratt 2008). We filmed emigrations with the camera positioned 70 cm above the arena. Filming ended when colonies stopped performing tandem runs, usually within 2 h after the destruction of the old nest.

We measured the body size of tandem leaders and followers by averaging three still images per ant taken from the video recordings (the correlation coefficients *r* among individual images was on average 0.93, i.e. measurement 1 vs. measurement 2 of the same ant, measurement 1 vs. measurement 3 of the same ant). The still images were taken at the beginning of a tandem run, so that the person taking the measurements was unaware whether a tandem run was going to be successful or not. Additionally, we measured the pair’s rate of progress (“speed”, cm/sec) and the walked distance by the pair (cm). To calculate the rate of progress, we subtracted the time ants were searching for each other during short interruptions from the total duration of the tandem run. This provides us with a tandem speed estimate that is unaffected by short interruptions. The walked distance is a measure of how direct tandem runs are. A tandem was considered successful (*i*) if the pair reached the nest entrance together or (*ii*) if the follower reached the nest entrance less than one minute after a contact loss. Only tandem runs from the old to the new nest were analyzed, *i.e.* forward tandem runs. Ants were not marked and to avoid measuring the same ant more than once (pseudoreplication), each pair was removed from the experiment immediately after the leader contacted the new nest. Ants that were part of unsuccessful tandem runs were also removed.

#### Part 4: body size and task

We measured the body size of five extranidal workers (presumably ants scouting for food or nest sites) and five intranidal workers (presumably nurses) from each of ten recently collected colonies. To make sure that some ants left the colony to explore the environment, colonies were starved for a few days. Extranidal workers were collected when they were encountered outside their artificial nest. Afterwards, the nest was opened and ants that sat on the brood pile or carried brood items were captured (intranidal workers). These ants were considered nursing workers. Each ant was photographed 3 times as described above.

### Statistical analysis

All statistical analyses were performed in R 3.4 (R Core Team 2016). For part 1, we quantified the worker size distribution and explored if body length showed significant skewness or kurtosis. For this we used the methods described by Crawley (2007, pp. 285-289). The values of each colony were first centered (mean = 0) and then combined to test overall patterns. To test if ant size affects walking speed (part 2), we used a general linear mixed-effects model (LME) with colony as random effect to control for the non-independence of data points from the same colony (Zuur et al. 2009). Size class was used as the fixed-effect. To analyze the effects of leader and follower body size on tandem running, both generalized and general linear mixed-effects models (GLMM’s and LME’s) were used. GLMM’s were used when the distribution of the response variable was binomial (success: yes or no), whereas LME’s were used to test the effects on speed, duration and distance. To test for the effect of body size variation on tandem running, we tested both the relative size difference (leader – follower, *i.e.* positive values mean that leaders were larger) and the absolute size difference (|leader − follower|, *i.e.* values always positive) as the predictors. To explore whether colony ID is important in explaining relationships, we (*i*) show results of both mixed-effects models and generalized linear models and (*ii*) compared mixed-models that include colony ID as a random effect with generalized linear models without a random effect. Likelihood ratio tests (LRT) were used to assess the significance of colony ID (Zuur et al. 2009).

## Results

### Part 1: body size distribution

Worker body length varied from 2.25 to 3.34 mm (2.68 ± 0.16, N = 262) in the four measured colonies (Fig. 1). Head width varied from 0.48 to 0.67 mm (0.58 ± 0.03). The within colony coefficient of variation (CV) for body length was 0.054 ± 0.01 (N = 4 colonies). Body size distribution was unimodal, suggesting that *T. nylanderi* does not have distinct physical castes (Fig. 1a). A Shapiro-Wilk test suggests a significant deviation from normality (W = 0.98472, *p* = 0.007). Body size also showed significant positive skew (*t*-value = 2.04, *p* = 0.02) and significant kurtosis (*t*-value = 3.55, *p* = 0.0002). The latter suggests significant pointiness (leptokurtic distribution).

**Figure 1.**
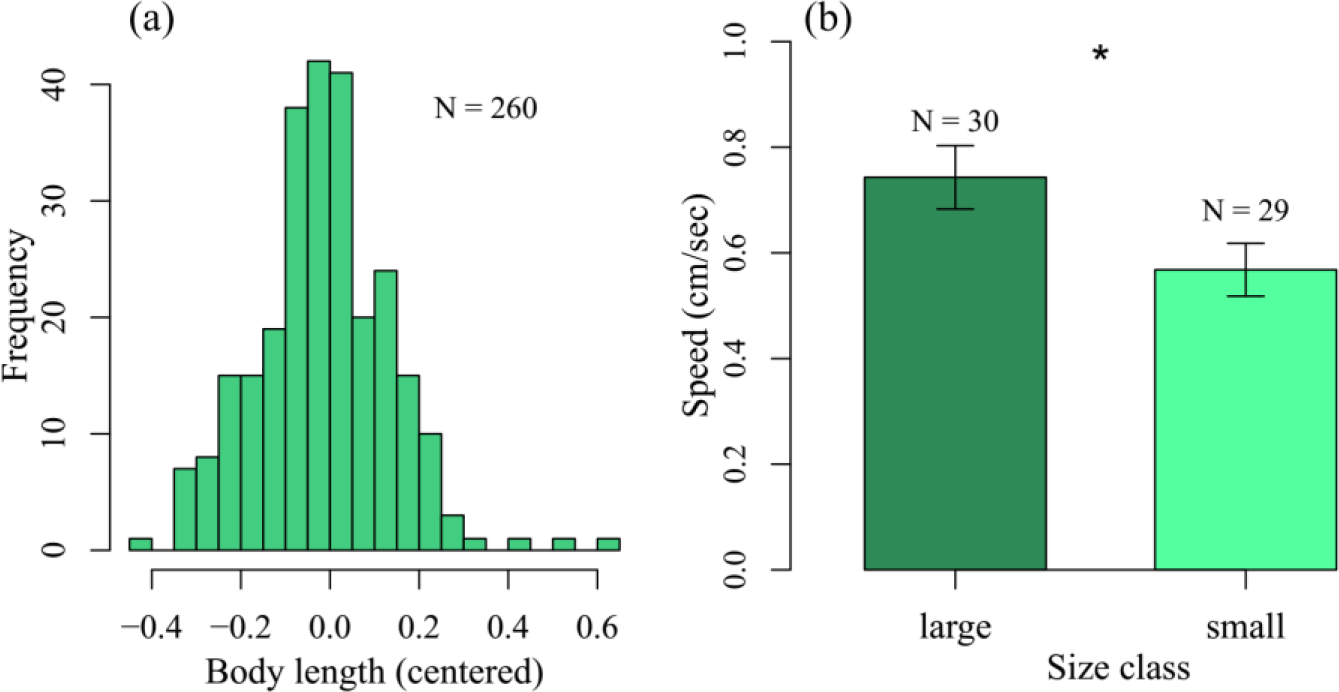
(a) Worker size (body length) distribution in *T. nylanderi* colonies. (b) The walking speed of individual workers depending on their size. Bars show means ± 1 standard error.

### Part 2: body size and walking speed

Ants from the large group were ~17% larger in body length than ants from the small group and walked significantly (+30%) faster than ants from the small group (Fig. 1b) (0.74 ± 0.3 cm/sec vs. 0.57 ± 0.26 cm/sec, LME: *t*-value = 2.32, *p* = 0.025).

### Part 3: Body size and tandem running

We analyzed 95 tandem runs from 22 colonies; 56% were successful. The size of tandem leaders did not correlate with the size of their followers (LME, values centered for each colony: *t*-value = −1.36, *p* = 0.18).

We then tested if the relative and absolute size difference predicted tandem success. Absolute size difference, but not relative size difference significantly affected tandem success (Fig. 2) (absolute difference: GLMM, z-value = −4.22, *p* < 0.0001; relative difference: z-value = −0.7, *p* = 0.49). Thus, larger size differences were associated with a low chance of tandem success, but it did not matter if the larger ant was the leader or the follower. We did not find any links between size difference and the speed, duration and walked distance of successful tandem runs (Table 1). We then explored whether the average size of ants in tandem (average of leader and follower) affects tandem success, but found no relationship (z-value = 1.31, *p* = 0.19). Generalized linear models provided nearly identical results (Table 1). The random effect colony ID was never significant (Table 1).

**Table 1.**
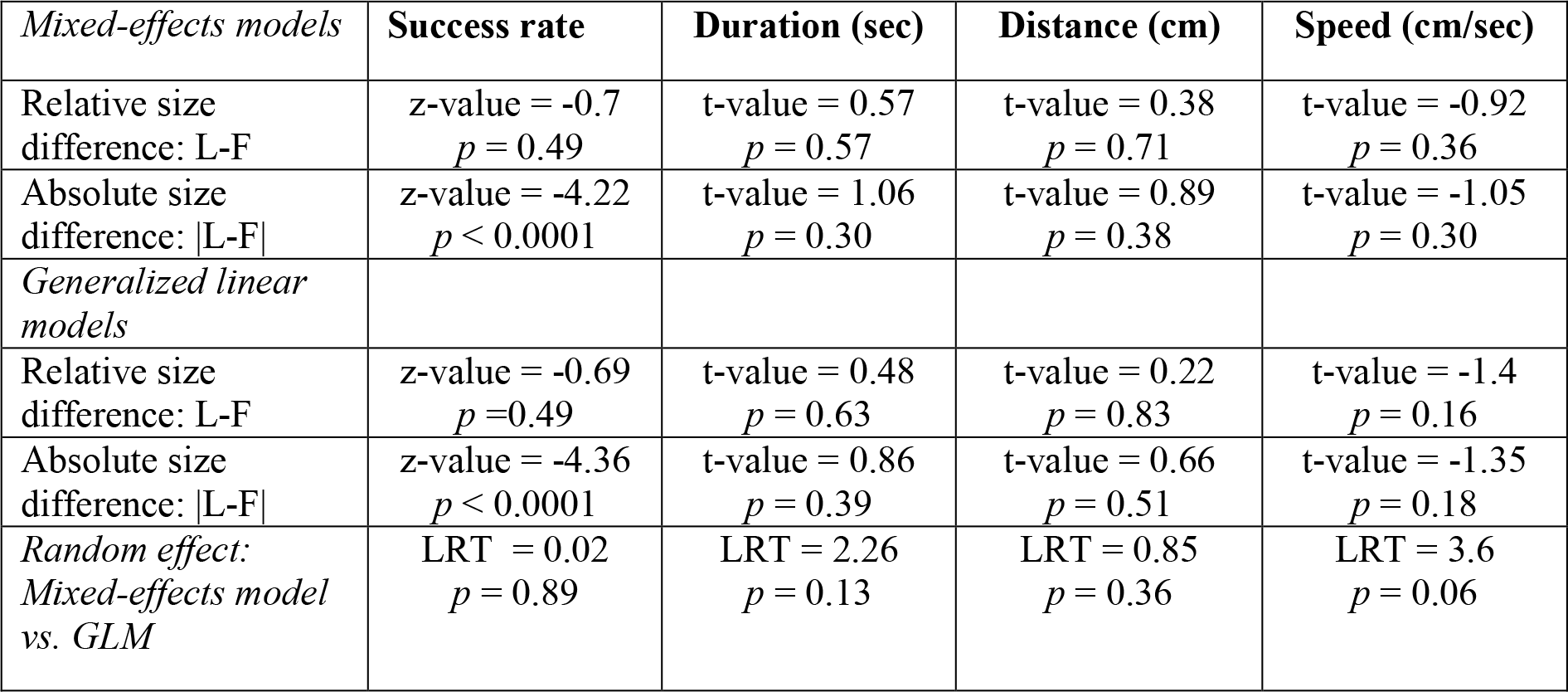
The effects of relative and absolute size difference on the speed (cm/sec), duration (sec, square root-transformed) and distance walked (cm, log-transformed) of successful tandem runs (N = 53). L = leader, F = follower. Linear mixed-effects models with colony as a random effect (GLMM’s and LME’s) and generalized linear models without random effect are shown (GLM’s). The two types of models were compared using a likelihood ratio test (LRT) to test for significance of the random effect.

**Figure 2.**
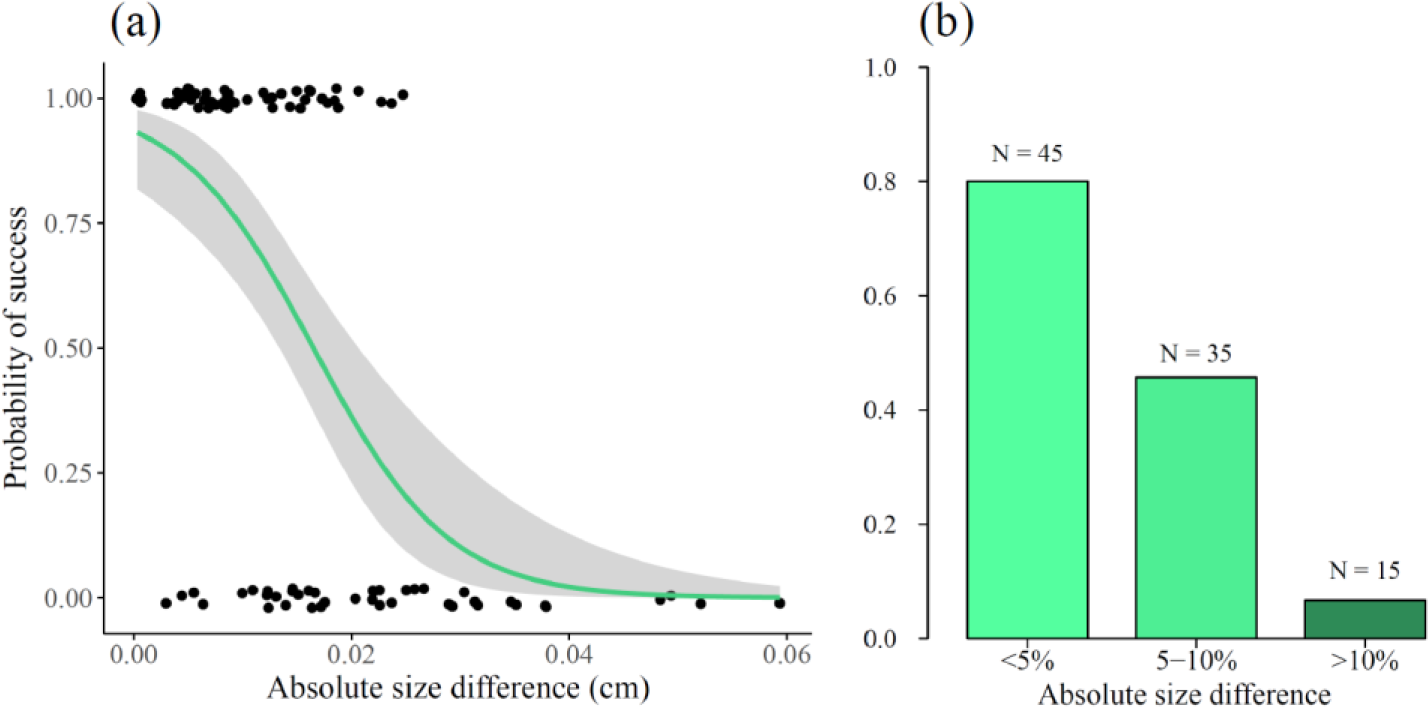
Probability that tandem runs were successful depending on the body length difference between leader and follower (a, b). (a) Note that all values are either 1 (success) or 0 (failure) but that jitter was used to better visualize the data points. Grey areas show the 95% confidence interval. (b) The same data as in (a), but the success probability for three different body length differences are shown. A value of 5% means that one of the ants was 5% smaller or larger than its partner.

### Part 4: body size and task

Extranidal workers (2.4 ± 0.2 mm; N = 50) were significantly larger than intranidal workers (2.32 ± 0.18 mm; N = 50) in ten recently collected colonies (LME: t-value = 2.12, p = 0.037).

## Discussion

We found that larger body size differences among interacting ants were associated with a high probability of tandem run failure. Only ~7% of tandem runs were successful when the leader-follower size difference exceeded 10%, whereas 80% of tandem runs were successful when ants differed less than 5% in body length (Fig. 2b). Speed, duration and distance travelled of the remaining successful tandem runs were not affected by body size differences. Our results suggest that size difference *per se* reduces tandem success, irrespective of whether the larger ant is the leader or the follower. One explanation could be that ants differing in size differ in their walking speed (Fig. 1b). Ants walking in a tandem run frequently need to accelerate and decelerate in order to maintain pair cohesion (Franks and Richardson 2006). Body size could affect the speed at which ants perform these changes and, thus, the probability of short contact losses. In this case, body size differences would not affect communication or signaling *per se*, but the ability of ants to stay together during this recruitment communication process. Alternatively, directional information that followers obtain from antennating the leader’s abdomen could be less precise if the partner differs greatly in size. It is also possible that the association between size variation and tandem success is driven by factors that were not measured in our study. For example, some colonies might be healthier or of generally better ability and this could affect both worker size variation and tandem running success. We would then predict that colony identity affects tandem success rate. However, we found no effect of colony identity on any of the measured parameters (Table 1).

Even though some lost followers may discover the new nest by themselves (Pratt 2008; Franks et al. 2010), unsuccessful tandem runs often represent a loss of time because lost followers require more time to discover a resource or they have to return to their nest following a break-up (Franks and Richardson 2006; Basari et al. 2014b; Grüter et al. 2018). Furthermore, a breakup could leave followers in unknown and dangerous locations. Alternatively, repeated partial tandem runs could represent a strategy to gain information about the direction to a new nest-site. However, given the search time of *T. nylanderi* followers after a break-up (Glaser and Grüter 2018) and the time ants have to wait inside the nest until they find a new leader, it is doubtful that this strategy would save time compared to performing one complete tandem run.

One observation that could indicate a strategy to reduce the risk of breakups is that extranidal workers were larger than intranidal workers. This is consistent with findings in two other *Temnothorax* species (Herbers & Cunningham 1983; Westling et al. 2014). Allocating larger workers to outside tasks is likely to decrease the probability that very large ants perform tandem runs with very small ants. Other potential advantages of larger extranidal workers could be an increased foraging rate due to their greater walking speed, greater foraging ranges (Ness et al. 2004, but see Westling et al. 2014) or a reduced risk of predation by certain predators.

Waddington (1989) hypothesized that size variation negatively affects waggle dance communication in honeybees because bees of different sizes may judge distances to food sources differentially. These costs of size variation could in turn select for low intra-colonial size variation (see also Waddington et al. 1986; Sauthier et al. 2017). While strong evidence for this is lacking, our finding that the size-frequency distribution in *T. nylanderi* colonies shows significant kurtosis (a leptokurtic distribution) indicates that there might be selection against body sizes that deviate strongly from the mean. Worker size variation in *T. nylanderi* (CV of ~ 5%) is similar to what has been found in *T. rugatulus*, which also performs tandem running (Westling et al. 2014). A comparison of worker size variation in species that use tandem running *versus* closely related species without tandem running could provide further evidence to support or challenge the hypothesis that the need for efficient recruitment communication selects for low intra-colonial worker size variation. Worker size in *T. nylanderi* is determined by food quantity, but also depends on rearing temperature and colony size (Molet et al. 2017). The degree of worker size variation could be modified either by nurse workers providing more constant or variable conditions (e.g. food amount) (Segers et al. 2015), or by larvae modifying their developmental response to external factors (Rissing 1987).

In *Apis mellifera*, dancing bees were more likely to be followed by bees of similar size (Waddington 1989). We found no correlation between leader and follower size in *T. nylanderi*. This could be explained by an inability of ants to accurately estimate the size of potential partners or by time costs that result from waiting for tandem partners of similar size. Such waiting time costs are likely to be larger in small colonies, such as in *T. nylanderi*, than in larger colonies, such as in honeybees (Anderson and Ratnieks 1999). Waggle dancing and tandem running are similar in that both require direct interactions between the signaler and the receiver. It is possible that behaviors involving direct interactions (e.g. cooperative transport, group recruitment or adult carrying) are more prone to being negatively affected by size variation than indirect forms of interactions, such as pheromone trails.

Body size variation is often linked to different roles in insect colonies. The most impressive examples are found in species with distinct physical worker castes (or sub-castes) performing different tasks (Wilson 1971; Oster and Wilson 1978; Hölldobler and Wilson 2009). However, most species display moderate, continuous worker size variation (Oster and Wilson 1978) and it is often not clear whether size variation has adaptive value at colony level. For example, bumblebees show moderate and continuous intra-colonial size variation (Goulson et al. 2002; Couvillon et al. 2010) and size is linked to the probability to perform certain tasks (Goulson et al. 2002; Spaethe and Weidenmüller 2002), but experimentally manipulating the body size variation in colonies did not affect colony performance in *Bombus impatiens* (Jandt and Dornhaus 2014). Similarly, a recent study using *T. nylanderi* found that experimentally reducing worker size variation did not affect colony performance in the laboratory for several tasks (Colin et al. 2017). This highlights that the effects of variation may not be straightforward and may remain hidden until colonies experience challenging conditions. Furthermore, while variation in some traits may be beneficial in the context of particular tasks, e.g. temperature regulation (Jones et al. 2004), variation in other traits or in the context of other tasks (e.g. communication tasks) may be inconsequential or even negative. In the latter case, colonies would have to trade-off the costs and benefits of variation and, as a result, measuring colony traits such as biomass or brood production would miss important consequences of worker diversity because it averages positive and negative effects. Another problem of colony level analysis is that natural selection might affect phenotypic variation only in particular sub-sets of colony members, e.g. the foragers. Future research can help identify the conditions and contexts that make variation an asset or a problem for insect societies and those where worker variation is simply the result of naturally occurring variation in environmental conditions, without having any fitness consequences for colonies.

## Acknowledgments

We thank Francisca Segers for comments on a previous version of the manuscript. S.G. and C.G. were funded by the Deutsche Forschungsgemeinschaft (DFG) (GR 4986/1-1).

